# Molecular detection of a cryptic salamander: Development of an eDNA assay for the detection of the mud salamander (*Pseudotriton montanus*)

**DOI:** 10.1101/2025.01.12.632641

**Authors:** Ben F. Brammell, Sara A. Brewer, Ethan A. Hoogerheide, Mary R. Johnson

## Abstract

The mud salamander (*Pseudotriton montanus*) is a notoriously cryptic semi-aquatic Plethodontid found throughout much of the eastern United States, reports of decades passing between observations of this species in areas of known occurrence are common. Although it is listed as imperiled or in need of conservation throughout much of its range, with extirpation suspected in many areas, relatively little is known of its current distribution due to its secretive nature. We developed a species-specific qPCR assay for use in eDNA detection of *Pseudotriton montanus*. Primers and probe were designed based on cytochrome b sequences obtained from specimens collected in central and eastern KY, compared to published sequences throughout the species’ range, and screened *in silico* (twenty-seven species) and *in vitro* (seventeen species) for specificity against sympatric salamander species. The developed assay was field tested via the collection of water samples at sites known or suspected to contain *P*. *montanus* in Kentucky, Ohio, and Tennessee. Of the 68 samples collected, *P. montanus* eDNA was detected in eight, including all sites (six) in which *P. montanus* larvae were observed in the field. Sequencing of each environmentally-obtained amplicon confirmed detection of *P. montanus diastictus*. This work provides thoroughly vetted tools that should prove useful for future monitoring and range delineation of this threatened and cryptic species.

## Introduction

Environmental DNA (eDNA) is increasingly utilized in ecological studies to address questions such as endangered species presence, expansion of introduced species (LaBrie et al. 2024; Tsuji et al. 2024), the spread of pathogens, and community composition evaluation. eDNA studies are most frequently conducted utilizing either PCR reactions that target a single species or high-throughput sequencing (metabarcoding) which enables rapid assessment of entire communities of taxa (Lodge 2022; Mu et al. 2024; Vanderpool et al. 2024). Although the relatively recent introduction of metabarcoding allows the detection of hundreds of species simultaneously, single-species assays may offer lower detection limits (Simmons et al. 2015; Harper et al. 2018; Bylemans et al. 2019; Wood et al. 2019; Banks et al. 2021; Moss et al. 2022), lower cost (Langlois et al. 2020), and are recommended over metabarcoding when study objectives are the detection of one or a small number of species (Goldberg et al. 2016; Wood et al. 2019). Single-species DNA assays remain an effective tool for the detection of species so rare that traditional survey approaches are either ineffective or prohibitive due to labor considerations (Wilcox et al. 2013; Duarte et al. 2023).

The mud salamander (*Pseudotriton montanus*, Baird 1849) is a relatively large (maximum TL = 21 cm [Petranka, 1998]), stout-bodied semi-aquatic plethodontid (Figure 1) found in the eastern United States from central Florida to New Jersey (Petranka 1998). *P. montanus* habitat has been broadly described as muddy areas associated with slow-moving streams and swamps in floodplain forests (Bruce 1975, 2003; Petranka 1998), although associations with headwater streams in areas lacking muddy substrates have also been reported (Hirschfeld and Collins 1963). The species has long been described as residing in low-elevation habitats (below 700 m) (Petranka 1998; Hunsinger 2005), with the Appalachian Mountains serving as a divider of east and west populations (Huheey and Stupka 1967), although recent studies report observations from higher-elevation habitats within the southern Appalachians (Hromada and Gienger 2020; Romans and Smith 2022). Breeding has been reported to occur in late summer to fall (Robinson and Reichard 1965; Bruce 1975) with oviposition occurring in fall or early winter in springs, seeps, and bogs (Brimley 1923; Fowler 1946; Goin 1947). Fully aquatic larval periods exceed one year (Bruce 1974, 1978), ensuring multiple larval cohorts at any given time (Goin 1951; Bruce 1978). Following metamorphosis, adults reside in or near water, often in association with fossorial burrows (Bruce 1975), or in nearby wooded areas under logs (Bruce 1975; Mount 1975).

**Figure 1.**
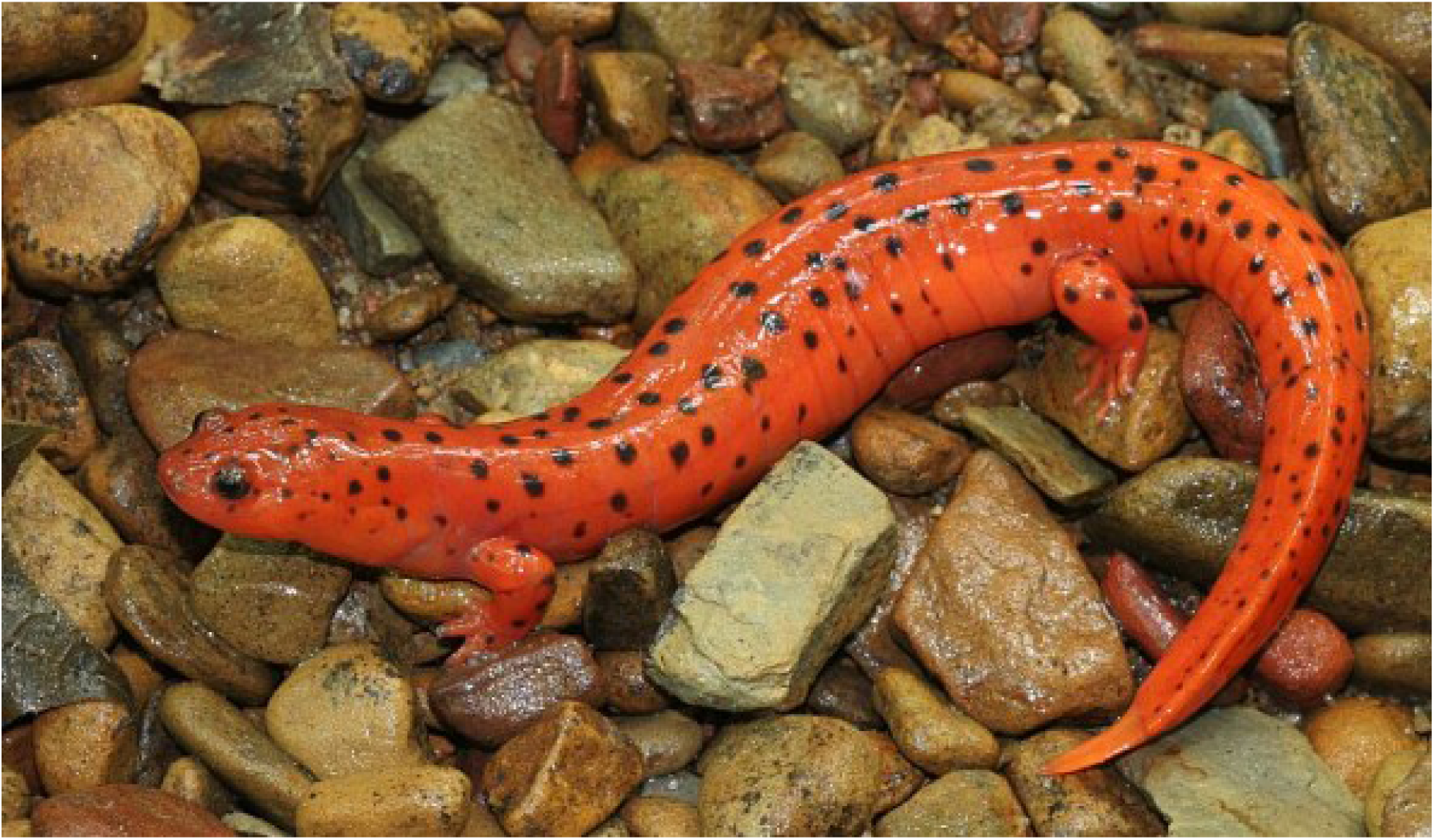
Large adult *Pseudotriton montanus* (mud salamander) observed in Vinton County, Ohio. Photo by Carl R. Brune.

*P. montanus* is notoriously cryptic, perhaps partially because of its subterranean tendencies, and little is known about the abundance of this species throughout much of its range (Petranka 1998; Hunsinger 2005). Reports of decades passing between observations in areas of known occurrence abound (McCoy 1992; Petranka 1998), and many accounts of this species are based on a single specimen or few individuals (Wright and Trapido 1940; Barbour 1953; Miller 1990). *P. montanus* was rediscovered in Louisiana in 1998 after a period of 31 years without a sighting (Hunsinger 2005) and in southern central Pennsylvania following an interlude of nearly 140 years (McCoy 1992). Of the sixteen states that comprise *P. montanus*’ range, this species is listed as a high priority conservation species in fourteen (Table S1), including several in which partial (KYDFWR 2023) or complete (Gessner and Stiles 2001) extirpation is suspected. While some state-listed statuses may be a result of the secretive nature of this species (Hunsinger 2005), improved monitoring is needed to fully understand the conservation status of this widely distributed but rarely observed species (Petranka 1998).

The life history and ecology of *P. montanus* make this species an ideal candidate for eDNA monitoring, and the development of eDNA protocols to detect this species has been listed as a conservation priority (KYDFWR 2023). The extreme cryptic nature of adults (Hunsinger 2005) and the fully aquatic larval phase lasting longer than one year (Bruce 1975), which guarantees that larvae will continually be in contact with the water if populations are present, present eDNA as a desirable option to include in the monitoring of this species. Additionally, larval *P. montanus*, particularly small larvae in the first few months of life, are difficult to distinguish from the frequently sympatric *Pseudotriton ruber* (Pfingsten and Brune 2013), which are often found in the same microhabitats (Pfingsten and Brune 2013). eDNA testing enables the differentiation of these two closely related species at any life stage. The objectives of the present study are to design and validate, *in silico*, *in vitro*, and *in situ*, a qPCR eDNA assay for use in the detection of *P. montanus*.

## Methods

### Tissue collection and sequencing

*P. montanus* tissue was obtained from tissue clips (< 2 mm) from specimens obtained in southeastern (2, Whitley Co., KY) and central Kentucky (1, Franklin Co., KY) (Table 1). Additionally, we collected tissue DNA from seventeen sympatric or potentially sympatric salamander species (Table 2) from central or eastern Kentucky for *in vitro* testing (KYDFW Permit #SC2111188). We extracted DNA from tissue using a DNeasy^®^ Blood & Tissue Kit (Qiagen) according to the provided protocol. Tissue was lysed overnight at 56°C in Proteinase K and eluted twice (200 µl and then 100 µl) to increase DNA yield.

**Table 1.** Comparison of *P. montanus* cytochrome b sequences currently (as of 8/31/24) found in GenBank and comparison of the similarity of these sequences. Sequences #1 through #3 were contributed by this study. The % similarity of each of the sequences are compared to Sequence #1.

| # | Subspecies | County | GB Acc. # | Len. (BP) | % similarity |
| --- | --- | --- | --- | --- | --- |
| 1 | *P. montanus diastictus* | Whitley Co., KY | MZ507697.1 | 839 | - |
| 2 | *P. montanus diastictus* | Whitley Co., KY | PQ197426 | 885 | 100 |
| 3 | *P. montanus diastictus* | Franklin Co., KY | PQ244009 | 893 | 98.7 |
| 4 | *P. montanus diastictus* | Lincoln Co., KY | KF562584.1 | 674 | 98.9 |
| 5 | *P. montanus diastictus* | Menifee Co., KY | KR054760.1 | 867 | 98.4 |
| 6 | *P. montanus montanus* | Wake Co., NC | KF562586.1 | 674 | 93.7 |
| 7 | *P. montanus flavissimus* | Macon Co., AL | KF562585.1 | 674 | 92.1 |
| 8 | *P. montanus flavissimus* | Macon Co., AL | KR054761.1 | 867 | 91.7 |
| 9 | *P. montanus floridanus* | Alachua Co., FL | MW319716.1 | 1,120 | 89.0 |

**Table 2.** Mismatches and amplification probabilities for *P. montanus* primers and probes and sympatric or potentially sympatric salamander species. “F+R primer” = forward and reverse primer, “P” = probe, “Amp. prob.” = probability of amplification of sequence with respective assay as determined by machine learning model (Kronenberger et al. 2022), “*In vitro”* = the primers were screened in laboratory tissue tests with this species.

| Sympatric species | Amp. prob. | F+R primer mismatches | P mismatches | GB Acc. # | *In vitro* |
| --- | --- | --- | --- | --- | --- |
| *Pseudotriton montanus* | 0.872 | 0 | 0 | MZ507697.1 | - |
| *Pseudotriton ruber* | 0.175 | 7 | 5 | MW319716.1 | Y |
| *Gyrinophilus porphyriticus* | 0.147 | 7 | 3 | KT794547 | Y |
| *Stereochilus marginatus* | 0.212 | 4 | 4 | JQ920617.1 | N |
| *Eurycea cirrigera* | 0.176 | 9 | 7 | JQ920622.1 | Y |
| *Eurycea lucifuga* | 0.131 | 6 | 5 | KT873718.1 | Y |
| *Eurycea longicauda* | 0.179 | 5 | 3 | AY528403.1 | Y |
| *Eurycea bislineata* | 0.222 | 3 | 5 | AY528402 | N |
| *Eurycea junaluska* | 0.198 | 7 | 3 | KF562550.1 | Y |
| *Eurycea wilderae* | 0.166 | 8 | 2 | PP393050.1 | Y |
| *Desmognathus fuscus* | 0.142 | 8 | 5 | MZ485476.1 | Y |
| *Desmognathus monticola* | 0.169 | 7 | 10 | MW319719.1 | Y |
| *Desmognathus ochrophaeus* | 0.097 | 9 | 9 | MW319718.1 | N |
| *Plethodon glutinosus* | 0.101 | 9 | 8 | MN723529.1 | N |
| *Plethodon dorsalis* | 0.174 | 11 | 7 | GQ464404 | N |
| *Plethodon richmondi* | 0.178 | 9 | 6 | AY378072 | N |
| *Ambystoma barbouri* | 0.151 | 8 | 6 | OL456142.1 | Y |
| *Ambystoma texanum* | 0.160 | 8 | 6 | OM236537.1 | Y |
| *Ambystoma opacum* | 0.095 | 8 | 6 | KT780868.1 | Y |
| *Ambystoma jeffersonianum* | 0.234 | 5 | 8 | KT780869.1 | Y |
| *Ambystoma maculatum* | 0.292 | 6 | 7 | EF036637.1 | Y |
| *Ambystoma tigrinum* | 0.208 | 5 | 7 | MZ962317.1 | Y |
| *Hemidactylium scutatum* | 0.152 | 8 | 8 | AY728231 | Y |
| *Notophthalmus viridescens* | 0.131 | 8 | 7 | AY691731 | Y |
| *Necturus maculosus* | 0.227 | 6 | 5 | KY788155.1 | N |
| *Cryptobranchus alleganiensis* | 0.115 | 8 | 7 | AY691719.1 | N |
| *Amphiuma means* | 0.238 | 3 | 6 | AY691722.1 | N |
| *Siren intermedia* | 0.097 | 8 | 5 | MH888034.1 | N |

### Sequencing of target species

We amplified a portion of cytochrome b (cytb) from *P. montanus* specimens (GenBank accessions #MZ507697, #PQ197426, #PQ244009) using primers which we designed (Table S2). Sequencing was conducted in duplicate by ACGT, Inc. (ACGTinc.com).

### Assay development and testing

We designed our assay based on the three cytb sequences we obtained from *P. montanus* specimens collected in eastern and central Kentucky. To exclude the possibility that our specimens were rare variants, we compared these three sequences to all other *P. montanus* cytb sequences found in GenBank (Table 1). Once we confirmed that the published *P. montanus* sequences were similar (89.0 – 98.9%) to our sequence, we designed a forward and reverse primer and probe (Table 3) using PrimerQuest™ software (IDT™) and aligned these with sympatric or potentially sympatric species using MEGA X and ClustalW to identify mismatches (Table 2). Twenty-seven sympatric or potentially sympatric salamander species were included in this alignment, including all interspecific associations throughout the *P. montanus* range (Hunsinger 2005) (with the exception *Desmognathus auriculatus* for which sequences were not available). GenBank accession numbers of the cytb sequences used in alignments are found in Table 2. Prediction of primer specificity based solely on the number and position of mismatches can be misleading (So et al. 2020); therefore, we used a machine learning program (eDNAssay software; Kronenberger et al. 2022) to predict probabilities of amplification in the same 26 potentially sympatric species (Table 2). Following confirmation of specificity of the assay, the forward and reverse primer as well as the probe containing a 5’ FAM reporter dye and 3’ ZEN/Iowa Black FQ quencher using PrimerQuest™ were ordered from Integrated DNA Technologies (IDT™; Coralville, IA, USA).

**Table 3.**
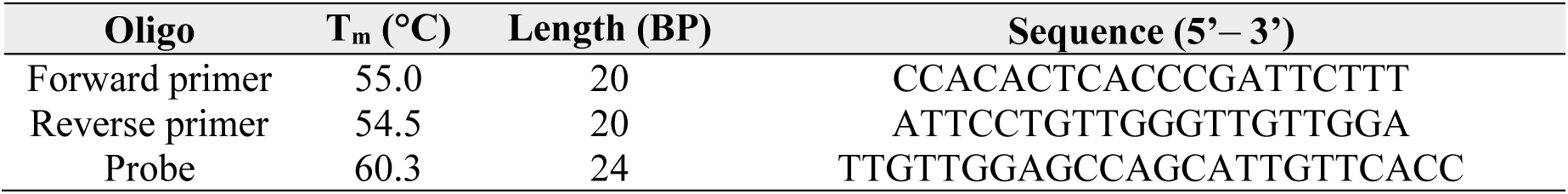
Quantitative PCR assay developed for *P. montanus*. The assay amplifies a 115 BP region of cytochrome b.

We evaluated the novel F and R primers via a polymerase chain reaction (PCR) temperature gradient approach to determine the optimal annealing temperature (nine temperatures between 57.4 – 65.9°C) of the primer set (Figure S1).

### *In vitro* specificity testing

For *in vitro* testing, we ran end-point PCR on tissue extracts of the six field-collected sympatric salamander species. Tissue-extracted DNA concentration was quantified using a Qubit™ 3.0 (Life Technologies, Carlsbad, CA, USA), and DNA from all species was diluted in nuclease-free water (IDT™) to a concentration of 1.0 µg/ml, exceeding concentrations used to verify *in vitro* specificity in previous studies (Witzel et al. 2020). Twenty-five µl reactions included: 12.5 µl GoTaq^®^ Master Mix (Promega), 8.5 µl nuclease-free water (IDT™), 2.0 µl tissue-extracted DNA, and 2.0 µl of F and R primers which had been diluted to 2 μM (final reaction concentration= 0.16 μM). PCR cycling conditions consisted of an initial denaturation stage of 95.0°C for two minutes followed by 35 cycles of 95.0°C for 60 s, 60°C for 60 s, and 72.0°C for 60 s, then a final extension of 72.0°C for five minutes. PCR products were visualized on agarose gels with 7.0 µl of PCR product loaded per well.

### Standard curves

We used synthetic DNA to assess assay sensitivity to enable standardization and interlaboratory comparison (Hobbs et al. 2019; Klymus et al. 2020). A 143-BP double-stranded synthetic DNA gBlock™ was ordered for *P. montanus* from IDT™. This consisted of the full eDNA primer-produced amplicon from our locally sequenced salamander DNA as well as additional bases which were added in order to exceed the minimum recommended length (Table S3).

We resuspended the gBlock™ in IDTE, pH 8.0 (IDT™), confirmed its concentration with a Qubit™ 3.0 fluorometer, and serially diluted it in nuclease-free water to produce a range of ten synthetic DNA concentrations from 10^7^ to 1 copies/μl. Two μl of each dilution were run in qPCR reactions with five technical replicates. The final range tested per reaction was therefore 2.0 x 10^7^ to 2 copies per reaction. Each 20.0 μl reaction contained the following: TaqMan™ EMM (10.0 μl), nuclease-free water (7.0 μl), resuspended gBlock™ (2.0 μl), and assay (1.0 μl). Thermocycler conditions were as follows: 50°C for 2 min, 95°C for 10 min then 55 cycles of 95°C for 15 s and 60°C for one minute.

The resulting data were plotted against copy number per reaction to determine the limit of detection (LOD) and limit of quantitation (LOQ). The LOD was defined as the lowest dilution of the standard curve that resulted in a detection of the target DNA with at least one qPCR replicate at a threshold cycle (Ct) of < 45 while LOQ was defined as the last standard dilution in which targeted DNA was detected and quantified in a minimum of 90% of qPCR replicates of the standard curve at a Ct < 45 (Mauvisseau et al. 2019).

### Field sampling and processing of eDNA samples

We collected stream water samples from locations in Kentucky, Ohio, and Tennessee from sites known or suspected to serve as *P*. *montanus* habitat in these states. Sample sites were selected based on historical data, herpetologist recommendations, and in two cases (including one of the positive detections) the density of sightings reported on the biodiversity social network “iNaturalist.” One-liter samples were collected in polyethylene containers that had not previously been exposed to the environment, containers were thrice rinsed in stream water before sample collection. Samples were transported on ice and either filtered or frozen within 48 hours. Water samples were filtered using vacuum filtration and 47 mm diameter glass microfiber filters (VWR, 0.42 mm thickness and 0.7 μm pore size) in a manner similar to previous studies (Jerde et al. 2011; Eichmiller et al. 2014; Guivas and Brammell 2020). Filter clogging is a frequently encountered issue in eDNA studies (Goldberg et al. 2016; Baldigo et al. 2017; Brammell et al. 2023), and one we observed in this study. We filtered as much water as possible from each habitat given the degree of clogging encountered. In 59/68 samples, one liter was filtered. For the remaining samples, greater than 600 ml was filtered for all except for three, in which case 500 ml, 350 ml, and 136 ml were filtered.

A comparison of field-based search methods with eDNA detection was not an objective in this study. As such, at field sites we only performed cursory field searches consisting of rock flipping and leaf litter sorting, depending on the environment encountered.

We extracted eDNA from filtered stream water samples using a modified version of a published protocol (Goldberg et al. 2011) for the DNeasy^®^ Blood & Tissue Kit (Qiagen), which has demonstrated superior yields relative to other extraction methods (Hinlo et al. 2017). Briefly, whole filters were cut into 10–12 pieces and incubated at 56°C overnight in 360 μl ATL buffer and 40.0 μl Proteinase K. Final elutions were performed twice (first with 200 μl and then with 100 μl of AE buffer).

We quantified environmental DNA using a StepOnePlus^TM^ Real-Time PCR system (Life Technologies™, Carlsbad, CA, USA) in optical eight-well PCR strips. Each run contained tissue-extracted target species DNA (1.0 μg/ml) as a positive control and also included a non-template negative control. Each 20.0 μl reaction contained the following: TaqMan™ EMM 2.0 (10.0 μl), nuclease-free water (2.0 μl), eDNA extract (7.0 μl), and assay (1.0 μl); primer and probe concentrations were identical to those previously described for the standard curve. Thermocycler conditions were as follows: 50°C for two min, 95°C for ten min, and 55 cycles of 95°C for 15 s and 60°C for one min. Each field-extracted sample was run in triplicate.

Inhibition testing was completed for field-extracted samples using TaqMan™ Exogenous Internal Positive Control Reagents (Applied Biosystems). Each 10.0 μl reaction contained the following: TaqMan™ EMM 2.0 (5.0 μl), nuclease-free water (0.3 μl), eDNA extract (3.5 μl), IPC assay (1.0 μl), and IPC DNA (0.2 μl). Thermocycler conditions were as follows: 50°C for two min, 95°C for ten min, 40 cycles of 95°C for 15 s, and 60°C for one min.

### Amplicon sequencing

Amplicons produced from environmental samples were sequenced, as recommended (Bockrath et al. 2022), to exclude the possibility of false positives. Amplicons were sequenced bidirectionally and therefore provided complete coverage of the entire sequence; all sequencing was conducted by ACGT, Inc. (ACGTinc.com).

## Results

### Cytochrome b sequencing

Of the three *P. montanus* cytb sequences obtained in this study, the two from specimens collected within the same county (Whitley Co., KY) are identical, and the third, collected approximately 160 km away in a more northern portion of the state (Franklin Co., KY), was 98.7% similar to these sequences (Table 1). Kentucky is home to only one *P. montanus* subspecies, *P. montanus diastictus* (Petranka 1998), and the two identical sequences from this study are greater than 98.4% similar to the two other *P. montanus diastictus* sequences available in GenBank (Table 1). Similarity of these two identical sequences with other *P. montanus* subspecies sequences in GenBank ranged between 93.7 and 89.0%, decreasing as the distance between the range of these subspecies and the range of *P. montanus diastictus* increased (Table 1).

### *In silico* specificity testing

The *P. montanus* oligos (F+R primer and probe) developed in this study (Table 3) have a minimum collective of seven mismatches with each sympatric salamander species considered (Table 3). Assay analysis with the eDNAssay machine learning software (Kronenberger et al. 2022) predicted an amplification probability of 0.872 for all three *P. montanus* cytb sequences acquired in this study while amplification probabilities of 0.292 or less were predicted for all non-target sympatric species cytb sequences (Table 2), meaning amplification would not be expected.

### *In vitro* specificity testing

End-point reactions with our *P.montanus* forward and reverse primers and six sympatric species revealed binding for target but not non-target species (Fig. S2). qPCR testing of our assay with tissue-extracted DNA likewise revealed amplification for target but not non-target species (Table 2).

### Assay validation

LOD and LOQ values were two and 10 copies per reaction, and the coefficient of determination (*r*^2^-value) for the curve was 0.98 with an amplification efficiency of 99.9% (Table S4).

### Field survey results

We considered a sample positive if at least one technical replicate out of three amplified (Harper et al. 2018; Plante et al. 2021). A total of eight samples among the 68 tested were positive for *P. montanus* DNA (Figure 2); details regarding the number of positive replicates and Ct values are found in Table S5). No amplification was observed in any of the field blanks.

**Figure 2.**
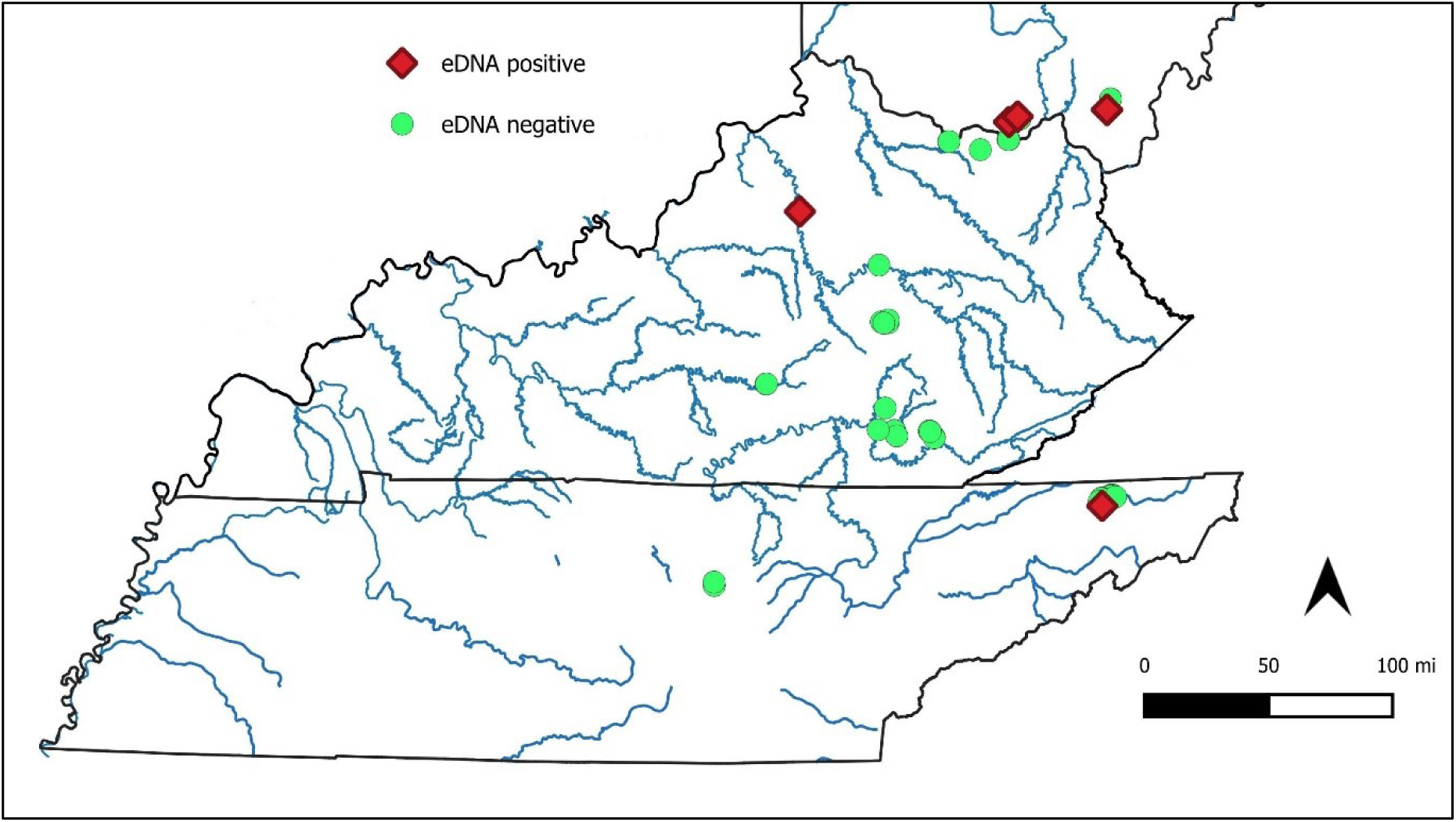
Water sample collection sites (68) for *P. montanus* eDNA detection. Green circles represent negative sites, red triangles represent sites positive for *P. montanus* eDNA. Samples too near one another appear as a single data point.

*P. montanus* larvae were observed at six of the 68 field sites. All larvae were photographed, and their identity was verified by at least one experienced herpetologist.

### Amplicon sequencing

Seven of the eight sequenced amplicons from the eight eDNA detections in this study were 100% matches to cytb sequences obtained in this study (GenBank accession #MZ507697.1 and #PQ197426) (Table S6). The last amplicon was 99.1% similar (1 base mismatch out of 115) to this sequence (Table S6).

## Discussion

The *P. montanus* eDNA assay designed in this study exhibits high sensitivity to low concentrations of target DNA (LOD ≤ 2 copies per reaction) and has been successfully validated *in silico*, *in vitro*, and *in situ*. Field validation led to positive detections at each site in which *P. montanus* larvae were observed (six) and at two locations in which no specimens were observed. *P. montanus* is uncommon throughout its range (Pfingsten and Brune 2013), with decades often elapsing between observations in areas of known occurrence (McCoy 1992; Hunsinger 2005). These observations are consistent with the results of this study, in which *P. montanus* was detected in a relatively low percentage of samples tested (8/68, 11.8), despite extensive efforts to select locations with histories of *P. montanus* occurrence. Site selection was made based on historic data, recent reports of adult sightings, and, in two cases, the density of “iNaturalist” sightings reported in a given area. Little is known regarding the abundance of *P. montanus* (Hunsinger 2005), and eDNA demonstrates great potential to serve as a valuable tool for future monitoring efforts of this cryptic species.

Intraspecific genetic variation presents a challenge to eDNA monitoring (Wilcox et al. 2015; Serrao et al. 2021; Brewer et al. 2024); in the case of *P. montanus*, its broad geographic range presents a unique opportunity to examine the effects of phylogeography on the efficacy of our eDNA assay. Although the taxonomy of *P. montanus* remains in need of study (Pfingsten and Brune 2013), four subspecies are currently recognized (Hunsinger 2005): the eastern mud salamander (*P. m. montanus*, found between the east coast of the U.S. and the Appalachians from Maryland southward to northern Georgia), the midland mud salamander (*P. m. diastictus*, which occupies a large area west of the Appalachians including much of Kentucky and Tennessee as well as portions of adjacent states), the Gulf Coast mud salamander (*P. m. flavissimus*, which is found from central Georgia in the coastal plain westward to eastern Louisiana), and the rusty mud salamander (*P. m. floridanus,* which is restricted to southern Georgia and the northern half of peninsular Florida (Bishop 1943)). Our assay was designed based on sequences obtained from specimens collected in southeastern and central Kentucky (GenBank accession numbers #MZ507697.1, #PQ197426, and #PQ244009), well within the range of *P. m. diastictus* (Bishop 1943). As expected, our assay demonstrated no mismatches with a published *P. m. diastictus* cytb sequence from a nearby southeastern Kentucky county (Lincoln Co., Cumberland River drainage) and only two mismatches with a published sequence from a specimen collected in eastern KY (Menifee Co., Kentucky River drainage) (Table S7). A comparison of assay effectiveness across subspecies reveals the expected effects of phylogeography; the available published sequences indicate our assay is effective for *P. m. diastictus* and *P. m. montanus*, but less effective for subspecies more geographically isolated from *P. m. diastictus* such as *P. m. flavissimus* and particularly *P. m. floridanus* (Table S7). However, given that *P. m. diastictus* and *P. m. montanus* comprise the majority of the *P. montanus* range, our assay should be broadly effective throughout the range of the species. Additionally, mismatched bases that could be altered to make our *P. montanus* assay a perfect match for the geographically isolated subspecies are presented in Table S7. Although *in vitro* testing would be required, these slight alterations to the oligos should render the assay completely effective in the detection of *P. m. flavissimus* and *P. m. floridanus*.

The development of accurate species-specific assays strongly depends on the availability of DNA sequences from closely related sympatric species (Duarte et al. 2023), and genetic variation due to phylogeography within these sympatric species can impede assay accuracy when utilizing it in increasing distances away from its location of origin (Ogata et al. 2022). In the case of *P. montanus*, its closest relatives are a monophyletic group consisting of two *Pseudotriton* species, *P. montanus* and *P. ruber*, *Stereochilus marginatus*, and members of the genus *Gyrinophilus*, consisting of the widely distributed *G. porphyriticus* and several endemic subterranean species. Both *P. ruber* and *G*. *porphyriticus* have extensive ranges in the eastern U.S. which largely overlap with *P. montanus*, and each are divided into four separate recognized subspecies (Folt et al. 2016; Kuchta et al. 2016). A comparison of the *P. montanus* assay presented here with published sequences from multiple specimens of each subspecies of *P. ruber* (Table S8) and *G. porphyriticus* (Table S9), as well as the three subterranean *Gyrinophilus* species (Table S9), indicates that our assay would be highly specific for *P. montanus* across the range of each of these sympatric species. Clearly, modification of our assay to be a perfect match to one of the geographically isolated *P. montanus* subspecies would require novel *in silico* analysis, but given the observed differences in this assay versus each sympatric species, it appears highly likely that the assay would remain accurate when considering these sympatric species.

Assay sensitivities reported here (LOD = 2, LOQ = 10) are similar to or lower than LOD and LOQ values reported by other eDNA studies. Kaganer et al. (2022) reported 10 copies/reaction as both the LOD and LOQ for a four-toed salamander (*Hemidactylium scutatum*) assay while Bell et al. (2022) reported values of 2 (LOD) and 20 (LOQ) copies/reaction for assays detecting southern two-line (*Eurycea cirrigera*) and northern dusky (*Desmognathus fuscus*) salamanders. An LOD of 23 and an LOQ of 75 copies/reaction were reported for an assay detecting California tiger salamanders (*Ambystoma californiense*; Kieran et al. 2020), while LOD values of 1, 100, and 1,000 copies/reaction were reported for assays detecting European crayfish (King et al. 2022). Guan et al. (2023) reported LOD’s of 4 and 8 and LOQ’s of 32 for two assays designed to detect Ridgeway’s rail (*Rallus obsoletus*). The sensitivity of the assay presented here is consistent with current eDNA single-species assay standards and appears capable of detecting DNA towards the lower end of eDNA-detectable concentrations.

The development and validation of novel eDNA assays requires significant levels of labor and resources (Wilcox et al. 2020); assays that have been thoroughly screened and validated hold value that extends well beyond their study of origin and provide valuable resources for facilitating future monitoring studies (Xia et al. 2021). Additionally, recent works have highlighted the need for both thorough specificity testing and standardization of assay validation (Goldberg et al. 2016; Klymus et al. 2020; Loeza-Quintana et al. 2020) as well as careful evaluation of sympatric species DNA to lessen the likelihood of false positives (So et al. 2020; Duarte et al. 2023). The *P. montanus* assay presented here has been validated to level four on the Thalinger scale (Thalinger et al. 2021) and screened *in silico* with interspecific associations throughout the *P. montanus* range (Hunsinger 2005) (except for *D. auriculatus*), including subspecies of the two most closely related species. The assay presented here provides a ready-made tool for molecular monitoring of this cryptic species and should serve as a valuable resource to enhance detection and monitoring efforts.

## Acknowledgements

We are indebted to many scientists who generously shared information and advice regarding likely positive control field sites for *P. montanus*. We thank Brett Kuss (Univ. of the Cumberlands), John MacGregor (KY Dept. of Fish and Wildlife), Carl Brune (Ohio University), John Howard (Southern Ohio Environmental Services, LLC), Jenny Richards (Shawnee State Park), Anthony Brais (Stan Tech), Bob Culler (Bays Mountain Park), and Sayre Stejbach (Northern Kentucky University, Center for Environmental Restoration). In addition, we thank Carl Brune (Ohio University) and John Howard (Southern Ohio Environmental Services, LLC) who provided valuable assistance in the field. Salamander tissue was collected under Kentucky Department of Fish and Wildlife Service Permit #SC1811153 (Brammell), and water samples in Tennessee were collected under Tennessee Wildlife and Resources Agency Permit #6095 (Brammell). All research was conducted in full compliance with both institutional and federal IACUC guidelines.

## Author contributions

BB served as the PI of the lab from which this research originated; he obtained funding, participated and supervised all lab work, collected a number of field samples, and wrote the manuscript text, preparing all figures and tables. SB obtained funding, conducted nearly all of the lab work for this study, collected many of the field samples, and contributed significantly to the writing of the manuscript. EH and MJ coordinated the locations of many field sites in the study and collected a number of field samples for the study. All authors reviewed and contributed to the manuscript.

## Declarations

The authors have no relevant financial or non-financial interests to disclose.

