## Supplementary Information for "Molecular detection of a cryptic salamander: Development of an eDNA assay for the detection of the mud salamander (*Pseudotriton montanus*)"

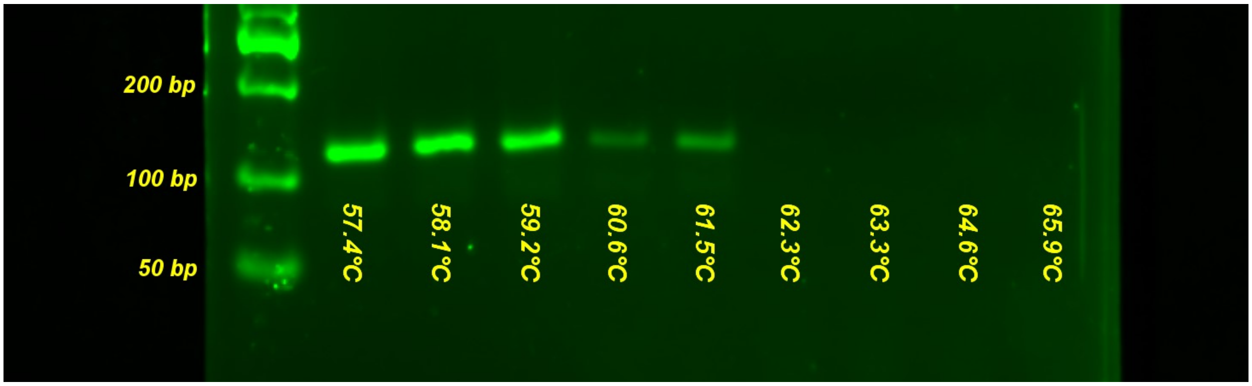


Figure S1. Gradient reaction to determine the optimal annealing temperature for the designed *P. montanus* primer set. Twenty-five µl reactions included: 12.5 µl GoTaq® Master Mix (Promega), 8.5 µl nuclease-free water (IDT™), 2.0 µl tissue-extracted DNA, and 2.0 µl of F and R primers. Cycling conditions consisted of an initial denaturation stage of 95.0°C for two minutes followed by 40 cycles of 95.0°C for 60 s, varying annealing temperatures for 60 s, and 72.0°C for 60 s followed by a final extension of 72.0°C for five minutes. PCR products were resolved on agarose gels with 7.0 µl of PCR product loaded per well.

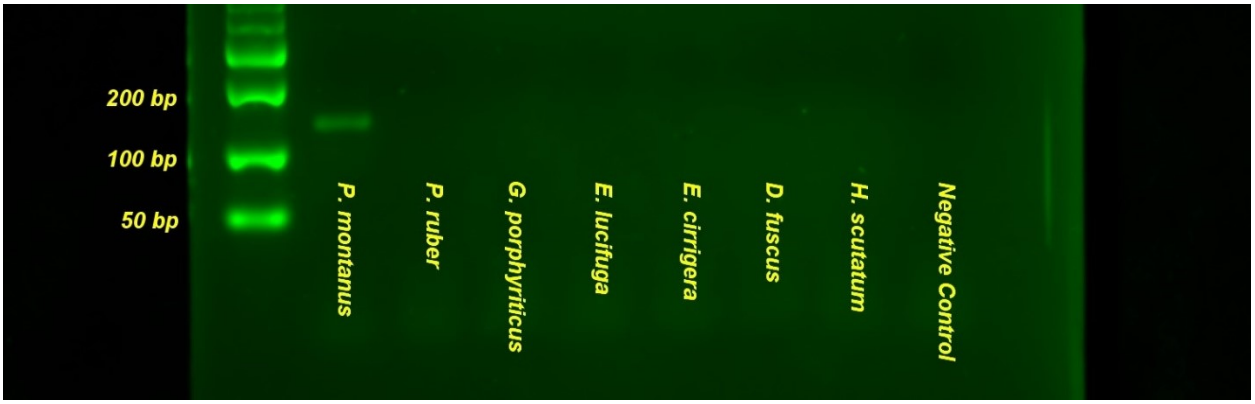


**Figure S2**. Tissue-extracted DNA specificity tests for *P. montanus*. End-point PCR reactions were conducted on tissue extracts of target species and six sympatric salamander species; all tissue was quantified (Qubit) and diluted to 1.0 µg/ml. Twenty-five µl reactions included: 12.5 µl GoTaq® Master Mix (Promega), 8.5 µl nuclease-free water (IDT™), 2.0 µl tissue-extracted DNA, and 2.0 µl of F and R primers. Cycling conditions consisted of an initial denaturation stage of 95.0°C for two minutes followed by 35 cycles of 95.0°C for 60 s, 60.0°C for 60 s, and 72.0°C for 60 s followed by a final extension of 72.0°C for five minutes. PCR products were resolved on agarose gels with 7.0 µl of PCR product loaded per well.

### Supplemental Tables

**Table S1.** Conservation status of *P. montanus* by state in each of the states included within the species range in the United States.

| State | *P. montanus* status by state | Reference |
|---|---|---|
| New Jersey | State threatened, possibly extirpated | (Gessner & Stiles, 2001) |
| Delaware | Endangered | (Delaware Division of Fish and Wildlife, 2013) |
| Maryland | Species of Greatest Conservation Need | (Chalmers & Therres, 2007) |
| Pennsylvania | Endangered | (Pennsylvania Fish and Boat Commission, 2024) |
| Ohio | Threatened | (Ohio Department of Natural Resources, 2022) |
| West Virginia | Species in Greatest Need of Conservation | (West Virginia Division of Natural Resources, 2015) |
| Virginia | Species of Greatest Conservation Need-Tier 4a | (Virginia Department of Game and Inland Fisheries, 2015) |
| Kentucky | Data deficient, possibly extirpated throughout much of its range in KY. | (Kentucky Department of Fish and Wildlife Resources, 2023) |
| North Carolina | Knowledge Gap, Research Priority | (N.C. Wildlife Resources Commission, 2015) |
| Tennessee | In greatest need of conservation | (Tennessee Wildlife Resources Agency, 2015) |
| South Carolina | Species of Conservation Concern, *Pseudotriton montanus flavissimus* | (South Carolina Department of Natural Resources, 2015) |
| Georgia | Not listed | (Georgia Department of Natural Resources, 2015) |
| Alabama | Moderate Conservation Concern | (Alabama Department of Conservation and Natural Resources, 2015) |
| Mississippi | Rare and/or imperiled | (Mississippi Commission on Wildlife, 2015) |
| Louisiana | Critically imperiled in Louisiana because of extreme rarity (5 or fewer known extant populations) or because of some factor(s) making it especially vulnerable to extirpation | (Department of Wildlife and Fisheries, 2015) |
| Florida | Not listed | (Florida Fish and Wildlife Conservation Commission, 2019) |

*LISTED SPECIES WILDLIFE THAT ARE CONSIDERED TO BE ENDANGERED, THREATENED, SPECIES OF CONCERN, SPECIAL INTEREST, EXTIRPATED, OR EXTINCT IN OHIO, Publication 5356*.

Pennsylvania Fish and Boat Commission. (2024, June 24). *Conservation, Salamanders*.

South Carolina Department of Natural Resources. (2015). *South Carolina's State Wildlife Action Plan*.

Tennessee Wildlife Resources Agency. (2015). *Tennessee State Wildlife Action Plan*.

**Table S2**. Primer sequences used in amplification of *P. montanus* cytochrome b for sequencing.

| GB Acc. # | Len. (BP) | Primers |
|---|---|---|
| MZ507697.1 | 957 | N. red F, Set #2 – CCAGCTCCGTCAAACCTATC<br>N. red R, Set #2 – GTCCTCCAATTCAGGTAAGTACAA |
| PQ197426<br>PQ244009 | 974 | Eurycea Cytb F – ACTCCTTTATTGACCTCCC<br>N. red R, Set #2 – GTCCTCCAATTCAGGTAAGTACAA |

**Table S3**. gBlock™ sequences used in qPCR assay sensitivity testing for *P. montanus*. Black bases represent amplicons (excluding the F and R primers), blue bases represent the F + R primers, pink bases represent probes, and red bases represent extra bases added to exceed the minimum length requirement (125 BP).

| Species | Total Length (BP) | Target Amplicon (BP) | gBlock™ Sequence |
|---|---|---|---|
| *P. montanus* | 143 | 115 | AAGTCAGCCTGCTCCACACTCACCCGATTCTTTGCTTTTC ACTTCATTTTACCATTTATAATTGTTGGAGCCAGCATTGT TCACCTATTATTTCTACATGAATCTGGATCCAACAACCC AACAGGAATACGCTGTGTCGTAAC |

**Table S4**. Limit of detection (LOD), limit of quantification (LOQ), coefficient of determination ($r^2$-value), and amplification efficiency (AE) for the designed qPCR assay.

| Species | LOD | LOQ | $r^2$ | AE |
|---|---|---|---|---|
| *P. montanus* | 2 copies per reaction | 10 copies per reaction | 0.9787 | 99.22% |

**Table S5.** Data for the eight samples positive for *P. montanus* eDNA.

| Sample | Date collected | Number of pos. reps. | CT value(s) |
|---|---|---|---|
| Franklin Co., KY #1 | 10/11/2023 | 3/3 | 37.5, 38.3, 39.4 |
| Sullivan Co., TN #2 | 1/4/2024 | 1/3 | 42.0 |
| Adams Co., OH #6 | 2/3/2024 | 1/3 | 40.0 |
| Scioto Co., OH #3 | 3/3/2024 | 1/3 | 40.2 |
| Scioto Co., OH #4 | 3/3/2024 | 1/3 | 41.3 |
| Lawrence Co., OH #1 | 3/17/2024 | 1/3 | 47.9 |
| Lawrence Co., OH #3 | 3/17/2024 | 3/3 | 38.3, 40.0, 45.0 |
| Lawrence Co., OH #4 | 3/17/2024 | 3/3 | 45.2, 47.0, 47.5 |

**Table S6.** Sequences of amplicons obtained from water samples. "% sim. to target species" represents a comparison of the amplicon with the cytochrome b sequences from *P. montanus diastictus* (GB Acc. #MZ507697.1 and #PQ197426) obtained in this study. Each amplicon was sequenced bidirectionally as unpurified PCR products to obtain the full amplicon (115 BP).

| Sample | Date collected | Length (BP) | % sim. to target species | Sequence |
|---|---|---|---|---|
| Franklin Co., KY #1 | 10/11/2023 | 115 | 100 | CCACACTCACCCGATTCTTTGCTTTTCACTTCATTTTACCATTTATAA TTGTTGGAGCCAGCATTGTTCACCTATTATTTCTACATGAATCTGGA TCCAACAACCCAACAGGAAT |
| Sullivan Co., TN #2 | 1/4/2024 | 115 | 100 | CCACACTCACCCGATTCTTTGCTTTTCACTTCATTTTACCATTTATAA TTGTTGGAGCCAGCATTGTTCACCTATTATTTCTACATGAATCTGGA TCCAACAACCCAACAGGAAT |
| Adams Co., OH #6 | 2/3/2024 | 115 | 100 | CCACACTCACCCGATTCTTTGCTTTTCACTTCATTTTACCATTTATAA TTGTTGGAGCCAGCATTGTTCACCTATTATTTCTACATGAATCTGGA TCCAACAACCCAACAGGAAT |
| Scioto Co., OH #3 | 3/3/2024 | 115 | 100 | CCACACTCACCCGATTCTTTGCTTTTCACTTCATTTTACCATTTATAA TTGTTGGAGCCAGCATTGTTCACCTATTATTTCTACATGAATCTGGA TCCAACAACCCAACAGGAAT |
| Scioto Co., OH #4 | 3/3/2024 | 115 | 100 | CCACACTCACCCGATTCTTTGCTTTTCACTTCATTTTACCATTTATAA TTGTTGGAGCCAGCATTGTTCACCTATTATTTCTACATGAATCTGGA TCCAACAACCCAACAGGAAT |
| Lawrence Co., OH #1 | 3/17/2024 | 115 | 100 | CCACACTCACCCGATTCTTTGCTTTTCACTTCATTTTACCATTTATAA TTGTTGGAGCCAGCATTGTTCACCTATTATTTCTACATGAATCTGGA TCCAACAACCCAACAGGAAT |
| Lawrence Co., OH #3 | 3/17/2024 | 115 | 100 | CCACACTCACCCGATTCTTTGCTTTTCACTTCATTTTACCATTTATAA TTGTTGGAGCCAGCATTGTTCACCTATTATTTCTACATGAATCTGGA TCCAACAACCCAACAGGAAT |
| Lawrence Co., OH #4 | 3/17/2024 | 115 | 99.1 | CCACACTCACCCGATTCTTTGCTTTTCACTTCATTTTACCATTTATAA TTGTTGGGGCCAGCATTGTTCACCTATTATTTCTACATGAATCTGGA TCCAACAACCCAACAGGAAT |

**Table S7.** Comparison of oligos developed in this study with the specimens of *P. montanus* found in GenBank. Numbers represent the number of mismatched bases between our oligos and the displayed sequence.

| Subspecies | County | GB Acc. # | Len. (BP) | Amp. prob. | F primer | R primer | Probe |
|---|---|---|---|---|---|---|---|
| *P. montanus diastictus*[1] | Whitley Co., KY | MZ507697.1 | 839 | 0.872 | 0 | 0 | 0 |
| *P. montanus diastictus*[1] | Whitley Co., KY | PQ197426 | 885 | 0.872 | 0 | 0 | 0 |
| *P. montanus diastictus*[1] | Franklin Co., KY | PQ244009 | 893 | 0.872 | 0 | 0 | 0 |
| *P. montanus diastictus* | Lincoln Co., KY | KF562584.1 | 674 | 0.872 | 0 | 0 | 0 |
| *P. montanus diastictus* | Menifee Co., KY | KR054760.1 | 867 | 0.758 | 0 | TCCAA~~C~~AACCCAACAGGAAT<br>T | TTGT~~T~~GGAGCCAGCATTGTTCACC<br>C |
| *P. montanus montanus* | Wake Co., NC | KF562586.1 | 674 | 0.761 | 0 | 0 | TT~~G~~TTGGAGCCAGCATTGT~~T~~CACC<br>A C |
| *P. montanus flavissimus* | Macon Co., AL | KF562585.1 | 674 | 0.631 | 1<br>C~~C~~ACACTCACCCGATTCTTT<br>T | 1<br>TC~~C~~AACAACCCAACAGGAAT<br>T | 3<br>T~~T~~~~G~~T~~T~~GGAGCCAGCATTGTTCACC<br>CA C |
| *P. montanus flavissimus* | Macon Co., AL | KR054761.1 | 867 | 0.631 | 0 | 1<br>TC~~C~~AACAACCCAACAGGAAT<br>T | 3<br>T~~T~~~~G~~T~~T~~GGAGCCAGCATTGTTCACC<br>CA C |
| *P. montanus floridanus* | Alachua Co., FL | MW319716.1 | 1,120 | 0.348 | 1<br>CCAC~~A~~CTCACCCGATTCTTT<br>C | 1<br>TC~~C~~AACAA~~C~~CCAACAGGAAT<br>T T | 4<br>TT~~G~~T~~T~~GG~~A~~GCCAGCATTGT~~T~~CACC<br>A C G C |

1 = Sequence obtained in this study.

**Table S8.** Comparison of oligos developed in this study with the four subspecies of *Pseudotriton ruber*. Numbers represent the number of mismatched bases between our oligos and the displayed sequence. With the exception of the *P. ruber ruber*, GB Acc. #OQ376719 (Brewer et al. 2024), all sequences, clades, and population numbers are from Folt et al. (2016).

| Subspecies | County | GB Acc. # | Clade | Pop. # | Amp. prob. | F+R primer mismatches | Probe mismatches |
|---|---|---|---|---|---|---|---|
| *P. r. ruber* | Madison Co., KY | OQ376719 | - | - | 0.175 | 7 | 5 |
| *P. r. ruber* | Rockcastle Co., KY | KR054853 | B4 | 41 | 0.175 | 7 | 5 |
| *P. r. ruber* | Menifee Co., KY | KR054854 | B4 | 42 | 0.175 | 7 | 5 |
| *P. r. ruber* | Clarke Co., GA | KR054858 | B4 | 19 | 0.175 | 7 | 5 |
| *P. r. ruber* | Athens Co., OH | KR054922 | B3 | 43 | 0.115 | 7 | 6 |
| *P. r. ruber* | Summit Co., OH | KR054897 | B3 | 45 | 0.115 | 7 | 6 |
| *P. r. ruber* | Moore Co., NC | KR054924 | B4 | 30 | 0.115 | 7 | 6 |
| *P. r. ruber* | Franklin Co., TN | KR054916 | B2 | 29 | 0.115 | 8 | 4 |
| *P. r. ruber* | Madison Co., AL | KR054845 | B2 | 23 | 0.115 | 8 | 4 |
| *P. r. nitidus* | Unicoi Co., TN | KR054871 | B4 | 38 | 0.175 | 7 | 5 |
| *P. r. nitidus* | Watauga Co., NC | KR054882 | B4 | 39 | 0.175 | 7 | 5 |
| *P. r. schencki* | Swain Co., NC | KR054880 | B4 | 36 | 0.175 | 7 | 5 |
| *P. r. schencki* | Graham Co., NC | KR054875 | B4 | 33 | 0.175 | 7 | 5 |
| *P. r. schencki* | Fannin Co., GA | KR054864 | B4 | 27 | 0.184 | 7 | 6 |
| *P. r. schencki* | Gilmer Co., GA | KR054859 | B4 | 24 | 0.184 | 7 | 6 |
| *P. r. vioscai* | Marshall Co., KY | KR054889 | A | 40 | 0.172 | 6 | 5 |
| *P. r. vioscai* | Winston Co., MS | KR054913 | A | 13 | 0.172 | 6 | 5 |
| *P. r. vioscai* | Washington Co., LA | KR054911 | A | 1 | 0.172 | 6 | 5 |
| *P. r. vioscai* | Covington Co., AL | KR054903 | A | 3 | 0.154 | 6 | 4 |
| *P. r. vioscai* | Burke Co., GA | KR054895 | B2 | 12 | 0.112 | 8 | 5 |

**Table S9.** Comparison of oligos developed in this study with the four subspecies of *Gyrinophilus porphyriticus* and with troglobitic *Gyrinophilus* species. Numbers represent the number of mismatched bases between our oligos and the displayed sequence.

| Subspecies | County | Coor. | GB Acc. # | Amp. prob. | F+R primer mismatches | Probe mismatches |
|---|---|---|---|---|---|---|
| *G. p. duryi*[1] | Powell Co., KY | 37.81722 -83.68083 | KT794547 | 0.147 | 7 | 3 |
| *G. p. duryi*[1] | Pike Co., KY | 37.24889 -82.49139 | KT794494 | 0.126 | 6 | 4 |
| *G. p. duryi*[1] | Pike Co., OH | 39.17052 -83.208389 | KT794506 | 0.126 | 6 | 4 |
| *G. p. porphyriticus*[1] | Indiana Co., PA | 40.42916-79.211111 | KT794443 | 0.126 | 6 | 4 |
| *G. p. porphyriticus*[2] | DeKalb Co., TN | N.A. | KY073017.1 | 0.153 | 8 | 3 |
| *G. p. porphyriticus*[2] | Cocke Co., TN | N.A. | KY073018.1 | 0.153 | 6 | 3 |
| *G. p. danielsi*[3] | Burke Co., NC | 35.62632-81.802003 | AY728230 | 0.146 | 8 | 3 |
| *G. p. danielsi*[4] | Yancey Co., NC | 35.75246-82.221697 | KF562576 | 0.136 | 7 | 3 |
| *G. p. danielsi*[4] | Wilkes Co., NC | N.A. | KF562577 | 0.146 | 8 | 3 |
| *G. p. dunni*[1] | Pickens Co., SC | 34.74222-82.843056 | KT794536 | 0.146 | 8 | 3 |
| *G. p. dunni*[4] | Oconee Co., SC | 34.92282-83.087761 | KF562578 | 0.146 | 8 | 3 |
| *G. p. dunni*[1] | Union Co., GA | 34.65222 -84.0325 | KT794542 | 0.146 | 7 | 3 |
| *G. subterraneus*[2] | Greenbrier Co., WV | N.A. | KY073019 | 0.126 | 6 | 4 |
| *G. subterraneus*[4] | Greenbrier Co., WV | N.A. | KF562582 | 0.126 | 6 | 4 |
| *G. gulolineatu*[4] | Roane Co., TN | 35.75972-84.451944 | KT794549 | 0.160 | 7 | 3 |
| *G. palleucus necturoides*[4] | Rutherford Co., TN | 35.93861-86.307222 | KT794555 | 0.147 | 7 | 3 |
| *G. palleucus palleucus*[1] | Jackson Co., AL | 34.70861 -86.18139 | KT794553 | 0.147 | 7 | 3 |

1 = From (Kuchta et al., 2016)
2 = From (Wray and Steppan, 2017)
3 = From (Mueller et al., 2004)
4 = From (Bonett et al., 2014)
